# Characterization of Nonmotor Symptoms in the MitoPark Mouse Model of Parkinson’s Disease

**DOI:** 10.1101/2020.11.02.365874

**Authors:** Monica R. Langley, Shivani Ghaisas, Bharathi N. Palanisamy, Muhammet Ay, Huajun Jin, Vellareddy Anantharam, Arthi Kanthasamy, Anumantha G Kanthasamy

**Affiliations:** Parkinson Disorders Research Program, Iowa Center for Advanced Neurotoxicology, Department of Biomedical Sciences, Iowa State University, Ames, IA 50011

**Keywords:** Parkinson’s disease, behavior, nonmotor, MitoPark

## Abstract

Mitochondrial dysfunction has been implicated as a key player in the pathogenesis of Parkinson’s disease (PD). The MitoPark mouse, a transgenic mitochondrial impairment model developed by specific inactivation of TFAM in dopaminergic neurons, spontaneously exhibits progressive motor deficits and neurodegeneration, recapitulating several features of PD. Since non-motor symptoms are now recognized as important features of the prodromal stage of PD, we monitored the clinically relevant motor and nonmotor symptoms from ages 8-24 wks in MitoPark mice and their littermate controls. As expected, motor deficits in MitoPark mice began around 12-14 wks and became severe by 16-24 wks. Interestingly, male MitoPark mice showed spatial memory deficits before female mice, beginning at 8 wks and becoming most severe at 16 wks, as determined by Morris water maze. When compared to age-matched control mice, MitoPark mice exhibited olfactory deficits in novel and social scent tests as early as 10-12 wks. MitoPark mice between 16-24 wks spent more time immobile in forced swim and tail suspension tests, and made fewer entries into open arms of the elevated plus maze, indicating a depressive and anxiety-like phenotype, respectively. Importantly, depressive behavior as determined by immobility in forced swim test was reversible by antidepressant treatment with desipramine. Collectively, our results indicate that MitoPark mice progressively exhibit deficits in cognitive learning and memory, olfactory discrimination, and anxiety-and depression-like behaviors. Thus, MitoPark mice can serve as an invaluable model for studying motor and non-motor symptoms in addition to studying pathology in PD.

## Introduction

Parkinson’s disease (PD) is a chronic, progressive neurodegenerative disorder affecting about 5 million people worldwide. The neuropathology of this disease is characterized by a loss of dopaminergic neurons in the substantia nigra (SN) of the brain, leading to a functional loss of dopamine in the striatum and severe motor deficits. Additionally, accumulations of abnormal alpha-synuclein (αSyn) proteins form Lewy bodies and Lewy neurites, both pathologic hallmarks of PD. Several genes have been linked with PD including *PINK1, Parkin, DJ-1*, and *LRRK2*; however, the vast majority of PD cases are considered idiopathic, implicating an etiologic role of environmental factors such as metals, pesticides, and other toxins in the development of the disease. Cardinal motor symptoms such as bradykinesia, tremor, rigidity, and postural instability are still classically used for the clinical diagnosis of PD. Neuroinflammation, oxidative stress and mitochondrial dysfunction are thought to contribute to the neurodegenerative processes of this disease ^1–3^. Current therapies, including levodopa (L-DOPA), monoamine oxidase inhibitors, and dopamine agonists, treat the symptoms yet ultimately cannot interrupt or slow down the neurodegenerative process. Furthermore, most commonly prescribed treatments do not address the full scope of symptomology in PD patients.

In addition to the characteristic motor symptoms, nonmotor symptoms such as hyposmia, sleep disturbances, gastrointestinal (GI) dysfunction, autonomic and cognitive deficits negatively affect the quality of life and cost of living for PD patients. Although often overlooked, nonmotor symptoms are a frequent cause of hospitalization and diminished quality of life for PD patients ^4^. Neuropsychiatric symptoms in PD include depression, dementia, anxiety, apathy, and cognitive dysfunction. Olfactory deficits are observed in more than 95% of those affected by PD, and depression is estimated to affect more than one-third of PD patients ^4–6^. Interestingly, a recent cross-sectional observational study found that nonmotor deficits precede motor symptoms in later-onset PD, while younger patients showed the opposite trend ^7^. Although behavioral tests are available to study various nonmotor symptoms in rodent species, large data gaps still remain in understanding the nonmotor phenotype of many toxin-based and genetic models of PD ^8^.

In many toxin-based models of PD, rodent species either do not suffer from PD-related nonmotor symptomology or their complete behavioral phenotyping has not yet been performed. For example, animals receiving intraperitoneal injections of MPTP do not display any olfactory impairment, although intranasal MPTP administration can functionally damage the olfactory epithelium ^9^. Although gastric emptying and small intestine transit are unaffected by MPTP, the toxin-induced loss of enteric dopaminergic neurons increases colon motility^10^. Several studies using rodent models reported that exposure to the neurotoxic pesticide paraquat or paraquat/maneb co-administration only induced anxiety- and depression-like behaviors ^11–13^. Interestingly, a recent study revealed that rotenone-treated zebrafish display motor, olfactory, and neuropsychiatric symptoms ^14^.

In transgenic mouse models, only a few αSyn mutant animal models reportedly show olfactory and GI functional changes ^15–17^. Parkin knockout mice have spatial memory impairments, but do not show evidence of olfactory dysfunction, anxiety, depression, or motor deficits ^18^. *Dranka et. al*. recently identified olfactory dysfunction in the LRRK2^R1441G^ mouse model ^19^. The presence of nonmotor symptoms in PD models would be particularly useful if they can be characterized as early onset and progressive similar to clinical PD. It is imperative that we develop and characterize models that recapitulate a broad range of nonmotor abnormalities. These models could then be utilized in the development of therapies to treat nonmotor symptoms and also to screen for adverse effects resulting from dopaminergic therapies.

The MitoPark mouse model, which recapitulates many of the hallmark features of PD, was created by selectively inactivating the mitochondrial transcription factor A (TFAM) in the nigrostriatal pathway, creating a conditional knockout driven by the dopamine transporter (DAT) promoter. MitoPark mice exhibit adult-onset progressive dopaminergic neurodegeneration, protein aggregation in nigral tissues, and L-dopa-responsive motor deficits ^20,21^. More recently, MitoPark mice were discovered to display certain nonmotor symptoms such as all-light- or all-dark-induced circadian rhythm dysfunction and early cognitive deficits ^22,23^. In the present study, we systematically characterized the nonmotor behavioral phenotype of the MitoPark PD mouse model. The overarching hypothesis is that MitoPark mice display nonmotor symptoms characteristic of PD.

## Results

### Progressive Motor Deficits in MitoPark Mice

Previous studies have characterized motor deficits in MitoPark mice ^20^. In our laboratory, some female mice exhibit poor condition after age 24 wks. Therefore, we decided to sacrifice at 24 wks instead of the previously reported 40 wks ^20,21^. As expected, male and female MitoPark mice revealed decreased horizontal and vertical activities which progressively worsened over time, beginning at 12-14 wks of age (Fig. 1A-C). MitoPark mice also spent significantly less time on the RotaRod from 12 wks onward (Fig. 1D). No significant changes were observed for grip strength at any age between MitoPark mice and littermate controls (Fig. 1E), indicating forelimb neuromuscular function remained intact. Similar to previous reports, MitoPark mice showed a significant reduction in body weight at 20 wks for males and at 22 wks for females (Fig. 1F, Supplementary Fig. 1B). Males and females differed significantly for certain behavioral parameters and were therefore plotted separately as well (Supplementary Fig. 1A-D).

**Figure 1.**
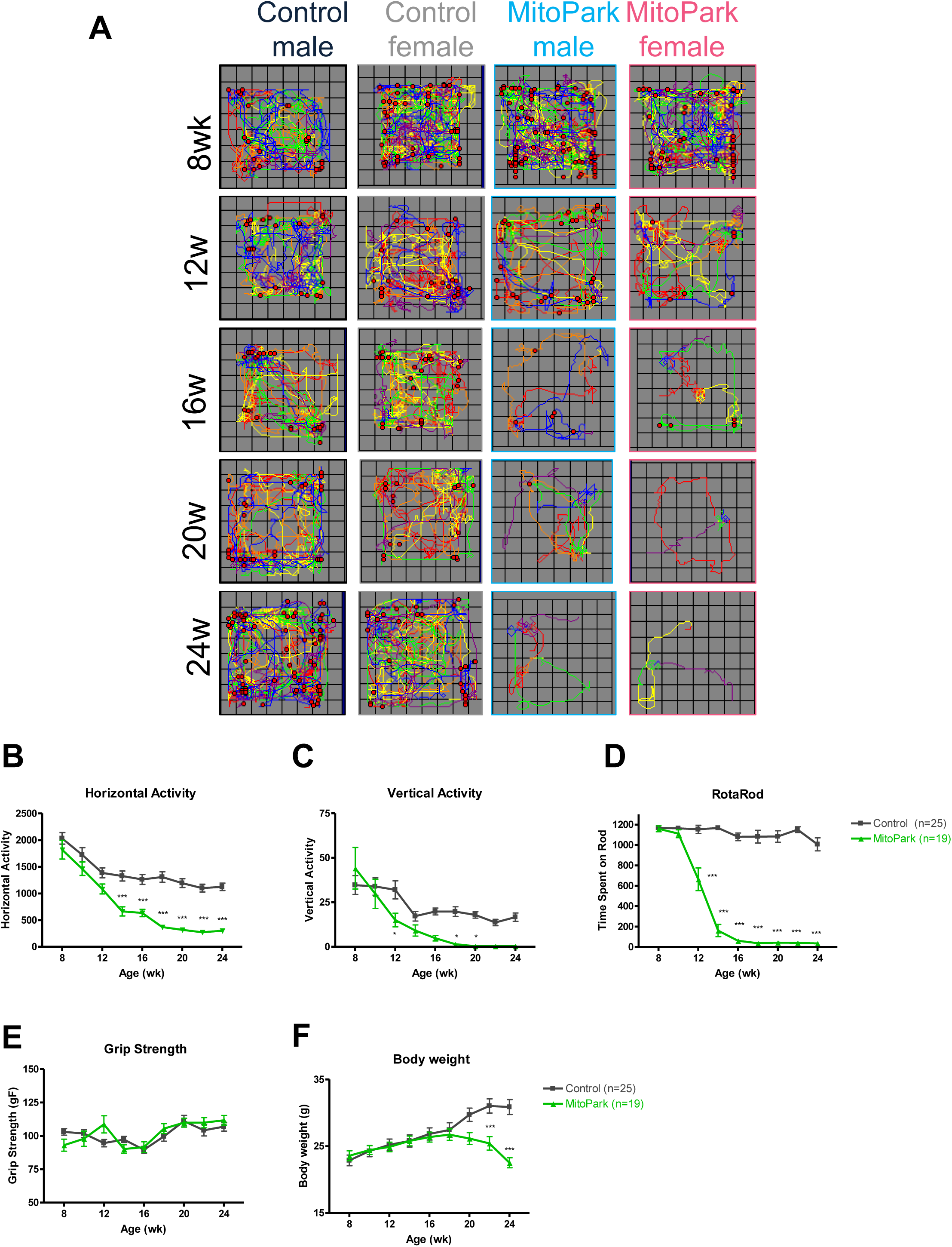
Progressive Motor Deficits in MitoPark Mice. A, Representative VersaPlots of individual mice subjected to the VersaMax open-field test showing horizontal (lines) and vertical (red dots) locomotion during a 10-min testing interval. Quantification of (B) horizontal and (C) vertical activities as determined by VersaMax analyzer during 10-min open-field test. D, Average time spent on RotaRod at 20 rpm during 5 trials. E, Grip strength and (F) body weights from 8- to 24-wk-old littermate control and MitoPark mice. *, p≤0.05; ***, p<0.001.

### Cognitive dysfunction in MitoPark Mice

To screen for cognitive deficits associated with spatial learning and memory, we next applied a six-day MWM protocol as described in Fig. 2A and depicted at 24 wks in Fig. 2B. Days 1-5 are track plots of an animal’s path to the visible (day1) or hidden (days 2-5) platform. Day 6 occupancy plots reveal time spent in each location during a one-minute retention trial with the platform removed. Importantly, average speed (m/s) was determined to not significantly differ between genotypes, and all animals were able to find the visible platform (day 1) at each age assessed (Supplemental Fig. 4). Recently, Li *et al*. ^23^ showed that cognitive dysfunction precedes motor deficits in MitoPark mice using the Barnes Maze. Similarly, we report that 8-wk MitoPark males exhibited impairments in the learning phase of the MWM (Fig. 2C). To our surprise, female MitoPark mice did not show an increase in escape latency until 12 wks of age (Fig. 2F-G). At 24 wks, both male and female MitoPark mice were unable to find the platform during the MWM learning phase (Fig. 2E, H). Deficits in the memory retention testing phase of MWM were apparent by 16 wks of age as depicted and quantified in Fig. 2I and 2J, respectively. Overall, these results not only confirm previous findings that learning deficits precede motor dysfunction in the MitoPark mouse model, but also further describe advanced spatial memory problems after 16 wks of age and reveal sex differences in learning the MWM platform location.

**Figure 2.**
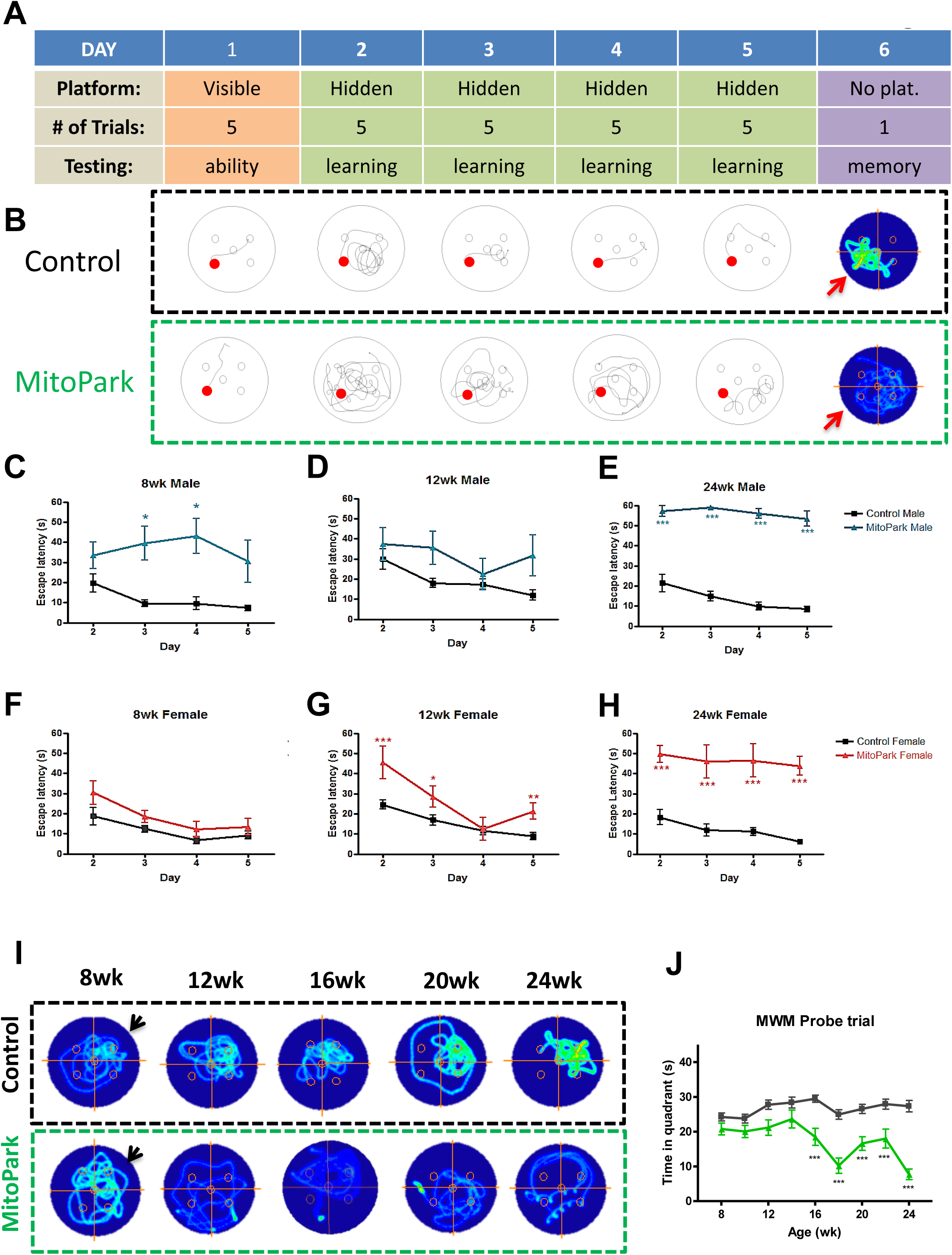
Cognitive dysfunction in MitoPark Mice. A, Schematic describing six day Morris water maze (MWM) protocol. B, Representative track plots from learning and occupancy plots from retention trials of MWM for one littermate control and MitoPark mouse. *C*, 8-, (*D*) 12-, and (*E*) 24-wk MWM learning period for male mice. *F*, 8-, (*G*) 12-, and (*H*) 24-wk MWM learning period for female mice. *I*, Representative occupancy plots of individual mice during retention trial with arrow indicating the previous platform location. *J*, Duration of retention trial spent in quadrant where platform had been located during learning phase. *, p≤0.05; **, p<0.01; ***, p<0.001.

### Neuropsychiatric symptoms in MitoPark mice

Depression is estimated to affect more than half of Parkinson’s patients and largely impacts patients’ quality of life ^24^. MitoPark mice were monitored every two weeks for depressive and anxiety-like symptoms from 8-24 wks of age. The tail suspension test (TST) revealed depressive-like behavior as indicated by increased immobility time in MitoPark mice at 16 weeks when compared to age-matched littermate control mice (Fig 3A). Control mice also showed increased immobility during TST at 24 wks. During the Porsolt forced swim test (FST), a significant increase in immobility occurred from 14 wks onward in MitoPark mice, while immobility in control mice remained relatively constant over time (Fig 3B).

**Figure 3.**
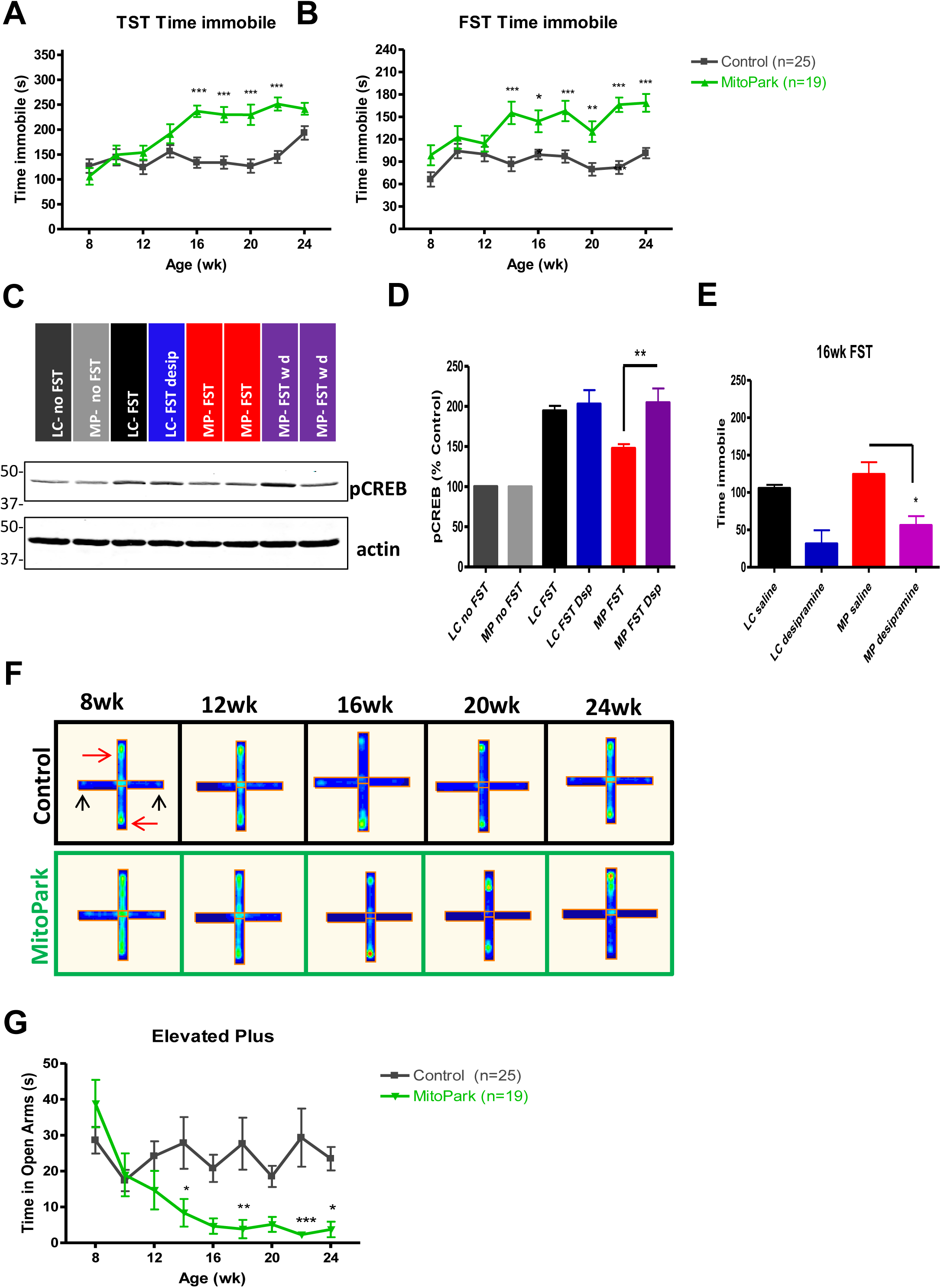
Neuropsychiatric symptoms in MitoPark mice. MitoPark mice were monitored at 2-wk intervals for neuropsychiatric deficits from 8-24 weeks of age. *A*, Tail suspension test (TST) and (*B*) forced swim test (FST) reveal depressive behavior in MitoPark mice at 16 and 14 weeks, respectively, when compared to age-matched control mice. *C-D*, Western blotting reveals increased CREB phosphorylation in FST-mice versus untested controls. *E*, Desipramine treatment (5 mg/kg, i.p., 30 min prior to FST) significantly reduced immobility time during the FST in MitoPark mice. *F*, Representative occupancy plots of closed (red arrow) and open (black arrow) arms of elevated plus maze and (*G*) corresponding time in open arms. *, p≤0.05; **, p<0.01; ***, p<0.001.

To further support that this finding was due to behavioral despair and not motor dysfunction, we treated a subset of 16- and 24-wk mice with desipramine (5 mg/kg, i.p.), an antidepressant that increases neurogenesis, and performed the FST 30 min post-treatment. In accordance with other studies showing antidepressant efficacy through neurogenesis, our Western blotting revealed increased CREB phosphorylation in the hippocampus of FST-tested mice versus untested controls (Fig. 3C-D). However, MitoPark mice did not show significant induction of pCREB unless treated with desipramine. Importantly, antidepressant treatment restored CREB phosphorylation to the levels in littermate control mice (Fig. 3C-D) and reduced immobility during the FST at 16 and 24 wks of age (Fig. 3E and Supplemental Fig. 3A). Male and female TST and FST data were plotted separately in Supplementary Fig. 1 D, and neurochemical data from desipramine-treated mice are available in Supplemental Fig. 3B. Our data show that increasing pCREB in the hippocampus of MitoPark mice attenuated behavioral despair. Furthermore, the fact that immobility was reduced by antidepressant treatment suggests that motor dysfunction in the MitoPark model is not the cause of immobility observed during the FST.

We also performed a 10-min elevated plus maze trial to test for anxiety-like behavior in MitoPark mice. The open arms are indicated by black arrows and closed arms by red arrows in Fig. 3F. Due to their natural preference for darker, enclosed spaces, mice in both groups spent less than 5-10% of their time exploring open arms. Despite the overwhelming preference for closed arms, clear behavioral differences emerged between groups. The time spent in the open arms decreased significantly in MitoPark mice beginning at 14 wks of age (Fig. 3G), suggesting a progressive increase in anxious behavior. Taken together, we have identified neuropsychiatric symptoms present starting from 14 wks in MitoPark mice, concurrent with the onset of motor deficits in this model.

### Olfactory dysfunction in MitoPark mice

Some degree of hyposmia is highly prevalent in PD patients and may occur decades prior to onset of motor dysfunction, making screening of olfactory deficits a potential prognostic tool in early PD ^25,26^. Representative occupancy plots from a 3-min trial of the social discrimination test (Fig. 4A) and novel scent test (Fig. 4B) at ages 8-24 wks reveal a reduced preference for the scented region (arrow) over time in MitoPark mice but not age-matched controls. Olfactory deficits, as indicated by significant differences in percent investigatory time during the social discrimination and novel scent tests, emerged at 14 and 16 wks, respectively (Fig. 4C-D). Also, fewer entries into the scented region during the social discrimination test occurred as early as 10 wks (Fig. 4E), while during the novel scent test, a significant reduction in entries began at 12 wks of age (Fig. 4F). Data is separated by sex in Supplementary Fig. 1 C. Our results indicate that olfactory deficits begin prior to the onset of motor symptoms in MitoPark mice.

**Figure 4.**
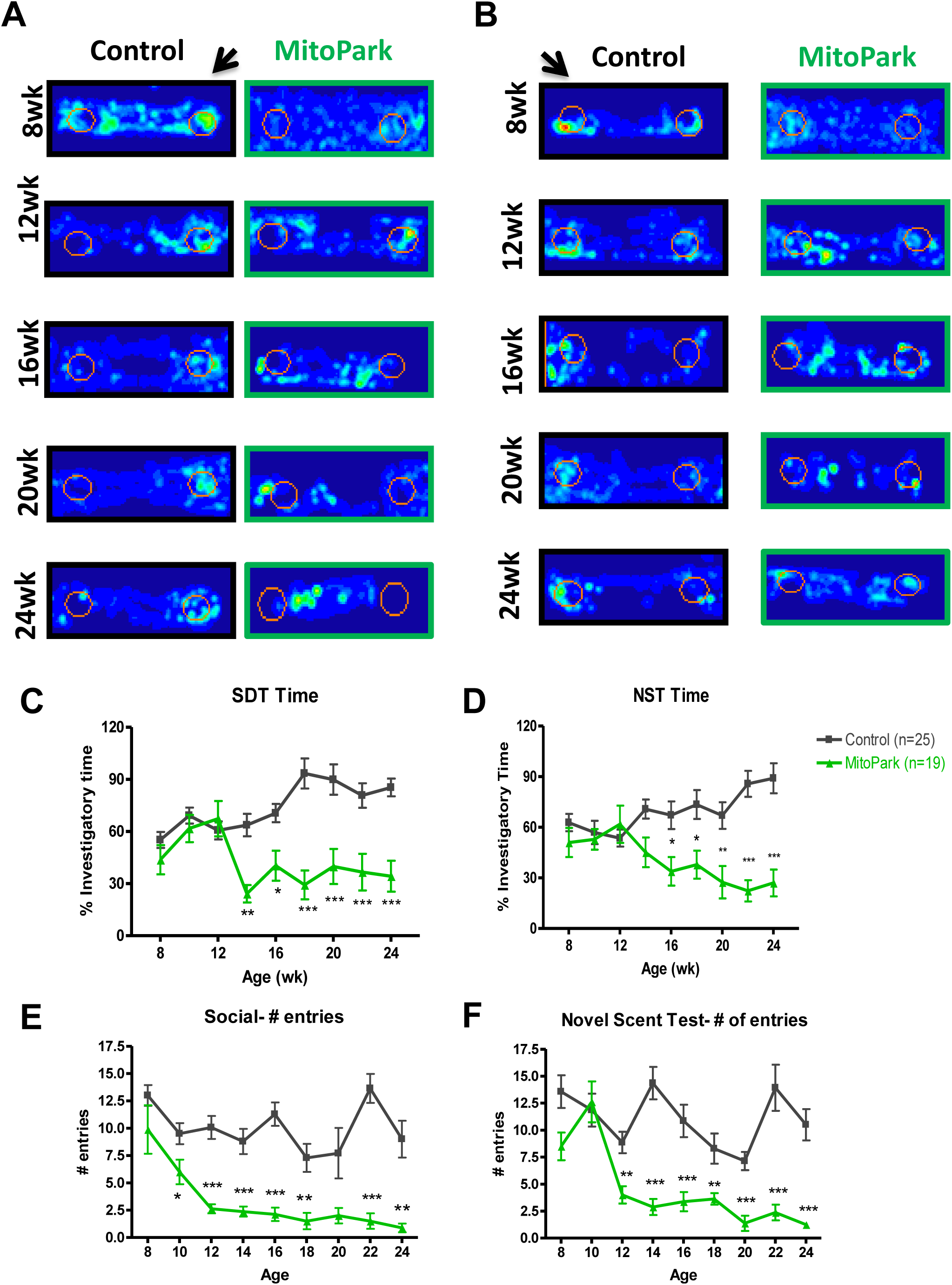
Olfactory dysfunction in MitoPark mice. MitoPark mice were monitored at 2-wk intervals for olfactory deficits from 8-24 weeks of age. *A-B*, Occupancy plots and quantification of (*C-F*) time in scented zones are shown. Olfactory deficits as determined by (*A, C, E*) social discrimination test (SDT) and (*B, D, F*) novel scent test (NST) were present as soon as 14 weeks of age in MitoPark mice versus controls. *, p≤0.05; **, p<0.01; ***, p<0.001.

### Biochemical changes correlating with the observed behavioral symptoms in MitoPark mice

Next, we tested for specific biochemical changes potentially corresponding to the occurrence of key non-motor symptoms. Since CREB phosphorylation and BDNF levels increase in mice post-MWM in a time-dependent manner^27–29^, we sacrificed mice 10-30 min after the last MWM retention trial and performed Western blotting for CREB, phospho-CREB, and BDNF protein levels. The significant reductions in CREB phosphorylation (Fig. 5A, C) and BDNF (Fig. 5A, D) protein levels in the hippocampus may be associated with the observed cognitive deficits.

**Figure 5.**
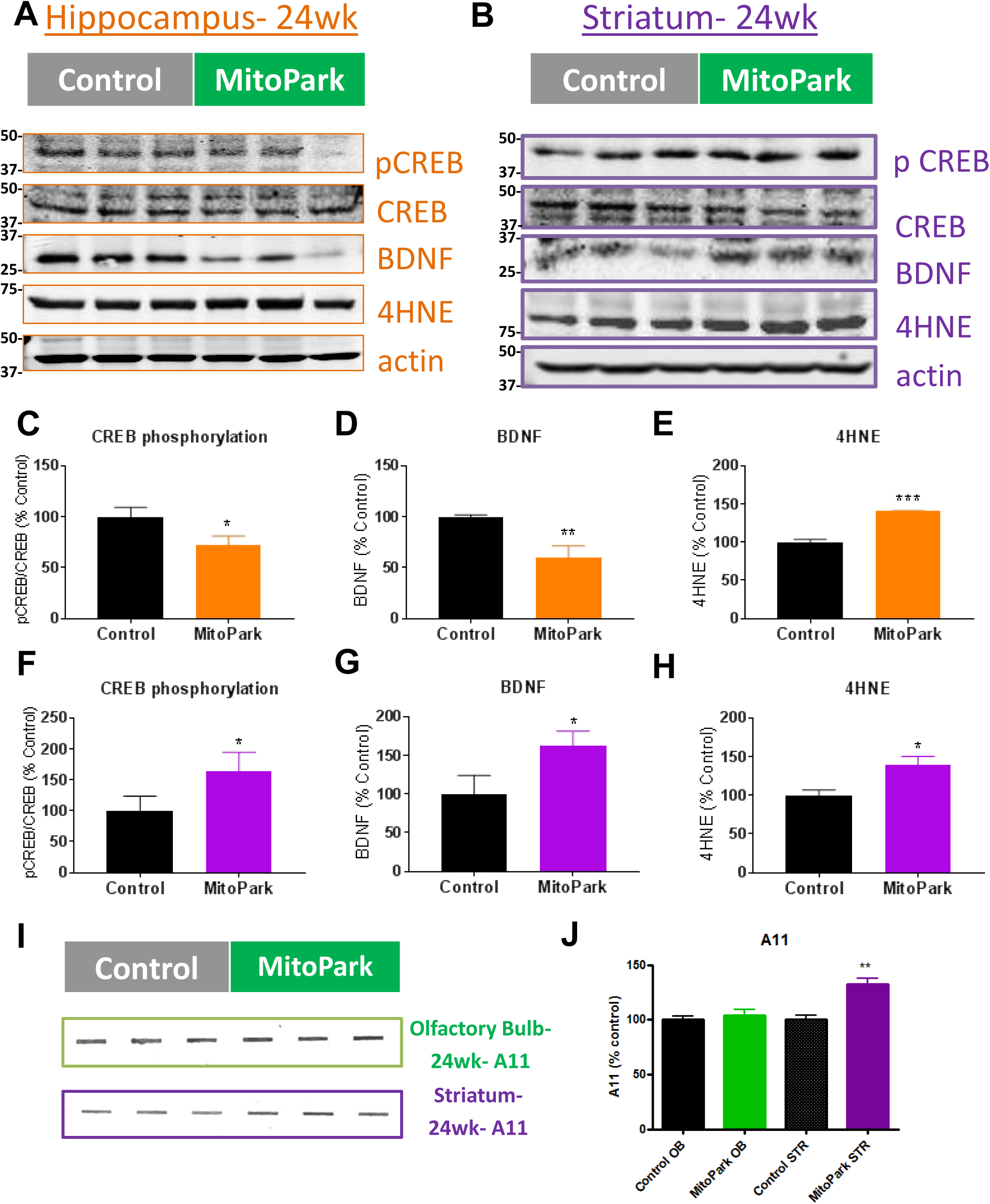
Biochemical changes correlating with observed behavioral symptoms in MitoPark Mice. *A-B*, Western blots and (*C-H*) densitometric analysis of proteins related to neuropsychiatric and cognitive changes. *I*, A11 slot blot and (*J*) densitometric analysis from striatum and olfactory bulb tissues from littermate control and MitoPark mice. *, p≤0.05; **, p<0.01; ***, p<0.001.

Conversely, the increases in CREB phosphorylation and BDNF in the striatum may be related to the depressive phenotype observed (Fig. 5B, F, and G). In both striatal and hippocampal tissues, we observed a significant increase in 4HNE, a lipid peroxidation product that results from oxidative damage (Fig. 5A, B, E, and H).

Researchers have attempted to link olfactory dysfunction to αSyn deposition, since both occur early in PD pathogenesis ^30^. However, we did not see an increase in oligomeric protein in the olfactory bulb of MitoPark mice as determined by slot blot for A11 anti-oligomeric protein antibody (Fig. 5I-J). Similar to what Ekstrand et al.^20^ reported in the substantia nigra of MitoPark mice, we did see significantly increased protein aggregation in striatal tissues.

### Neurochemical changes in MitoPark mice

Depression-like behaviors in toxin-based models of PD have been predominantly associated with reductions in hippocampal serotonin and striatal dopamine ^31^. However, no significant changes were observed in hippocampal neurotransmitters (Fig. 6A), indicating that observed changes in depressive behavior are instead possibly due to neurogenesis or plasticity modifications. As anticipated, strong reductions in dopamine and its metabolites were observed in the striatum of MitoPark mice, corresponding to the motor phenotype of the model (Fig. 6B). In our study, levels of dopamine and serotonin were reduced in the olfactory bulbs of 24-wk MitoPark mice (Fig. 6C), which may help explain the hyposmia in this model. However, *Branch et. al*. reported a dopamine reduction in the olfactory bulb that did not occur until a later age ^32^. Because DOPAC also increased (Fig. 6C), we believe enhanced dopamine turnover occurred in the olfactory bulbs of MitoPark mice. Since oligomeric protein did not significantly increase in the olfactory bulb (Fig. 5I-J), the hyposmia most likely resulted from neurochemical changes rather than protein aggregation. Future studies should explore the role of specific olfactory receptors in this model at various stages to better elucidate the mechanism of hyposmia in MitoPark mice.

**Figure 6.**
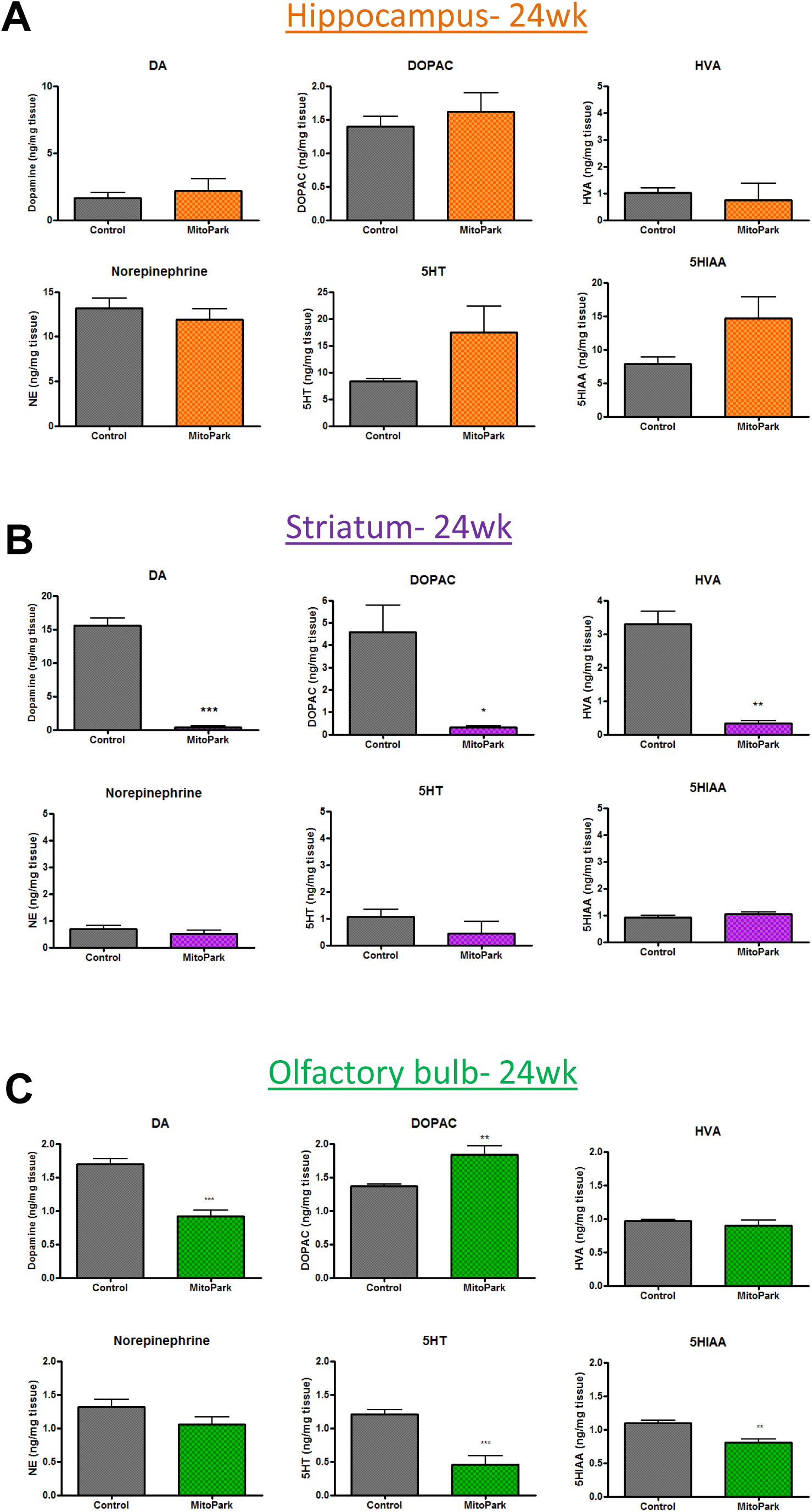
Neurochemical changes in MitoPark mice. Analysis of monoamine neurotransmitters from (*A*) hippocampus, (*B*) striatum, and (*C*) olfactory bulb tissues from MitoPark mice and their littermate controls at 24 wks of age. *, p≤0.05; **, p<0.01; ***, p<0.001.

## Discussion

Nonmotor symptoms often appear in the early stage of PD and can be more troublesome than the characteristic motor symptoms. Not only are they increasingly recognized as being critical to identify and treat due to their impact on the quality of life in PD patients, they could prove to be crucial prodromal indicators that could lead to better outcomes for PD patients by allowing treatments to begin sooner. MitoPark mice recapitulate several features of PD in humans, including a progressive course of the phenotypic manifestations and neurodegeneration, protein inclusions in nigral tissues, motor deficits that are ameliorated by L-DOPA administration, altered response to L-DOPA treatment, and adult-onset of disease. However, the full range of nonmotor symptomatology in the MitoPark model has not been characterized. In this study, we demonstrate that similar to human PD, many nonmotor symptoms including olfactory dysfunction, learning and memory deficits, and neuropsychological problems are also evident in MitoPark mice prior to or concurrent with the occurrence of motor deficits.

More than 70% of PD patients present nonmotor symptoms, according to a recent cross-sectional observational study^7^. Onset of the disease in areas of the brain outside of the substantia nigra (SN) or even peripheral onset of PD are supported by the idea that PD patients experience a variety of nonmotor symptoms before classical motor signs are observed ^33^. Certain symptoms are now considered to be early warning signs of PD, including hyposmia, constipation, rapid eye movement behavior disorder, and depression^34,35,36,37^. Although nonmotor symptoms typically occur prior to motor deficits in later-onset PD, recent studies suggest this is not true for those with young onset age of PD^7^. We found that while some behavioral symptoms in MitoPark mice occur prior to onset of motor symptoms (e.g., learning and olfactory deficits), many did not appear until after. For this reason, it is particularly important to consider parallel motor task parameters when interpreting the neurobehavioral results. For example, tasks which require horizontal movements (ie elevated plus, novel scent test) should be observed alongside open field test data which suggest motor deficits at 14 wks of age; whereas motor ability assessed during Morris Water Maze trials as average swimming speed (m/s) suggested no significant impairment in swimming. Interestingly, dual task performance has been shown in certain situations to improve performance for PD patients, thought to be due to increased production of catecholamines and arousal of additional brain regions ^38, 39^. Similarly, paradoxical kinesia can enhance motor performance during a perceived stressful event, such as an earthquake, and has also been observed to occur in PD^40^. These phenomena could potentially explain the enhanced ability of MitoPark mice to swim during Morris Water Maze during times when their locomotor activity is diminished in other types of motor tasks. Our cognitive function data are also supported by a recent study showing spatial learning and memory deficits in the Barnes maze and object recognition deficits that preceded the appearance of motor deficits in these animals ^22,23^. However, our study expanded their findings by showing various ages of mice, sex differences, and progressively worsening. Importantly, we have shown that MitoPark mice display clinically-relevant olfactory deficits, cognitive dysfunction, depressive- and anxiety-like behaviors that progressively worsen over time during adulthood.

The olfactory system provides sensory information from our environment that can be key to determining palatability of food and identifying the presence of dangerous fumes and other indicators of toxins^41^. The prevalence of hyposmia is about 90% in sporadic PD cases, and 68% of patients surveyed described alterations in their quality of life due to impaired olfactory function^41^. Olfactory disturbances do not occur in all neurodegenerative diseases, making this nonmotor symptom particularly valuable in differential diagnosis^42^. Dopaminergic cells can be found in the periglomerular cells of the olfactory bulb, but do no degenerate in PD ^41,43,44^. Unlike the SN and VTA, tyrosine hydroxylase (TH) actually increases in the olfactory bulb (OB) of patients with PD and in experimental PD animal models ^41,45,43,46,47^. In an intranasal MPTP model, dopaminergic and noradrenergic deficits were seen in the brain, but dopamine therapy does not help the olfactory deficits in PD patients^48^. Non-dopaminergic neurotransmitter systems are thought to contribute to or cause olfactory loss in PD^41^. The olfactory bulbs from older-aged MitoPark mice should be used to determine effects on protein aggregation and olfactory receptors. Changes in other components involved in olfactory discrimination should also be explored including odor receptors in the nasal epithelium, individual glomerular and mitral layers of the olfactory bulb, piriform cortex, hippocampus, and thalamus^49,50,51^.

Significant overlap and co-morbidity exists between nonmotor symptoms. For example, patients with hyposmia and rapid eye movement (REM) behavior disorder (RBD) are more likely to also exhibit cognitive deficits^37^. Both olfactory deficits and RBD were more prone to occur in patients with an older age of disease onset, and in such patients, these nonmotor symptoms were likely to occur prior to development of motor symptoms^7^. Combined effects or interdependency of behavioral phenotypes have not been addressed in this study.

Since the model was developed, more detailed pathological features have been identified by various groups. At 6-10 wks, impaired electrophysiological parameters were found in dopaminergic neurons, and increased L-type calcium channel mRNA and PK2 protein expression were observed in MitoPark mice versus their littermate controls ^52,32,53^. At 28-30 wks, increased astrocyte marker GFAP, greater striatal glutamate release, and white matter MRI changes were identified^54^. Also at 30 wks, MRI changes indicative of iron accumulation were found in the substantia nigra^54^.

We also observed strong neurochemical and biochemical changes that corresponded well with key non-motor symptoms. For example, reduced levels of dopamine and serotonin in the olfactory bulb of 24-wk MitoPark mice may help explain the hyposmia in this model ^55,56^. Within the mesolimbic dopamine circuit, increased BDNF through CREB activation mediates susceptibility to stress ^57,58^. In contrast, stress decreases hippocampal levels of BDNF and neurogenesis through CREB activity and cortisol concentrations ^57^. Oxidative stress has also been implicated in a variety of neuropsychiatric disorders, with particularly strong evidence in depression and anxiety studies ^59,60^. Also, reductions in CREB phosphorylation and BDNF in the hippocampus may be associated with the observed cognitive deficits and decreased neurogenesis. Conversely, increases in CREB phosphorylation, BDNF, and oxidative damage marker 4-HNE in the striatum may be related to the depression-like behavior observed. Although animal models and post-mortem studies do indicate adult neurogenesis is affected in PD, the exact mechanism of the changes and the correlation with nonmotor symptoms in this model have not been directly explored. Future studies should explore the role of adult neurogenesis in relation to nonmotor deficits in PD models.

Interestingly, no significant changes in sleep latency were observed (Fig. S2), although a recent paper found that MitoPark mice display circadian rhythm dysfunction following an all-light or all-dark cycle ^22^. Because constipation and other GI problems are associated with early stages of PD, further studies should be done to comprehensively characterize GI changes in MitoPark mice. Based on MPTP studies resulting in loss of dopaminergic neurons in the enteric nervous system, changes in colon motility should specifically be explored ^10^. Studies in our laboratory to comprehensively characterize GI function of MitoPark mice versus their littermate controls are currently underway (Ghaisas et al, 2018, unpublished).

Animal models and clinical PD studies have shown that reduced dopamine levels correlate with a reduced proliferation of cells in neurogenic regions of the brain, which is thought to contribute to the nonmotor symptoms observed in PD ^61–63^. However, for most mouse models of PD, either nonmotor symptoms do not emerge or else nonmotor performance was never characterized, presenting a narrow understanding of therapeutic potential with current genetic and toxin-based models ^8^. Interestingly, mice deficient in monoamine storage capacity display a progressive loss of dopaminergic cells in the SN, loss of striatal dopamine, motor deficits, αSyn accumulation and key nonmotor deficits^56^. Mitochondrial dysfunction is expected to lead to oxidative damage in neurodegenerative disorders ^64^, potentially representing the mechanism underlying the occurrence of neuropsychiatric symptoms in the MitoPark model. Neuroinflammation is another mechanism implicated in the pathophysiology of PD and nonmotor symptoms, given that inflammatory cytokine levels have been positively correlated with nonmotor deficits in a number of recent studies ^65,66^. We recently identified morphological changes indicative of microglia activation and increased IBA1^+^ microglia in the SN of MitoPark mice^67^.

Many nonmotor deficits closely correlate with Lewy body deposition and begin in the prodromal stage of PD, prior to the classical motor deficits used for clinical diagnosis ^68–70^. First, GI and olfactory disturbances are observed while αSyn pathology is evident in the olfactory bulb and the motor nucleus of the vagus nerve, which provides parasympathetic innervation to the GI tract ^71^. Next, synucleopathy in the hypothalamus, locus coeruleus, and raphe nucleus correlates with sleep disorders and neuropsychiatric symptoms such as depression and anxiety. Motor symptoms and clinical diagnosis then begin during Braak stages 3-4, when the midbrain becomes involved. Finally, cognitive decline and dementia are found in association with cortical deposition of αSyn, similar to what is seen in dementia with Lewy bodies ^72^. In addition to protein aggregate deposition, alterations in adult neurogenesis and neurochemical changes are implicated in the development of nonmotor symptoms in PD ^73,74^. Recent studies suggest that dopamine depletion and changes in αSyn may even synergistically contribute to the altered neurogenesis associated with nonmotor deficits ^73^. In our samples, we did observe increased striatal protein aggregation by anti-oligomeric (A11) slot blot analysis. Despite immunoreactivity to an anti-αSyn antibody, protein inclusions found in dopaminergic neurons of MitoPark mice were found not to contain αSyn ^20^.

Collectively, our study demonstrates that in addition to progressive motor deficits, MitoPark mice also exhibit nonmotor symptoms including olfactory dysfunction, learning and memory deficits, and neuropsychological problems. The presence of these symptoms in combination with the progressive motor dysfunction makes the MitoPark mouse model of PD particularly valuable for mechanistic and drug discovery studies.

## Materials and Methods

### Chemicals

Dopamine hydrochloride, 3-4-dihydroxyphenylacetic acid (DOPAC), and homovanillic acid (HVA) were all purchased from Sigma (St Louis, MO). Halt protease and phosphatase inhibitor cocktail was obtained from Thermo Fisher (Waltham, MA). Bradford assay reagent and Western blotting buffers were purchased from Bio-Rad (Hercules, CA). Anti-4-hydroxynonenal antibody was purchased from R&D Systems (MAB3249, Minneapolis, MN ^75^), while anti-BDNF was purchased from Santa Cruz Biotechnology (sc-546, Dallas, TX). CREB and p-CREB (Ser133) antibodies were obtained from Cell signaling (9104, 87G3, Boston, MA ^76^). The anti-mouse and anti-rabbit secondary antibodies (Alexa Fluor 680 conjugated anti-mouse IgG and IRdye 800 conjugated anti-rabbit IgG) were purchased from Invitrogen and Rockland Inc., respectively.

### Experimental design

Experimenter was blinded to mouse genotype for each behavioral test by flipping of cage cards and randomizing order prior to beginning the experiment. A third party was used to conceal treatment and vehicle solution identity prior to drug administration. Genotype and administration blinding was decoded after data acquisition was complete. Power analysis using the “fpower” function in SAS was used to determine the following animal requirements based on 80% power, alpha=0.05, and four groups. Clinically significant differences and standard deviations used to determine delta were taken from ANOVA analyses of previous studies in our lab. Based on preliminary forced swim test data, a minimum of 11 animals per group were used for behavioral animal studies to detect an effect size of 1.5. Male and female mice were combined, for a total of 19 MitoParks and 25 Littermate control mice used for all behavioral experiments. Mice unable to swim to a visible platform were to be excluded from Morris water maze on day one, however, no such mice were observed in our study.

### Animal treatment

MitoPark mice were originally kindly provided and generated by Dr. Nils-Goran Larson at the Karolinska Institute in Stockholm ^20^. All mice for this study were bred, maintained, genotyped, and further characterized at ISU. MitoPark mice (DAT +/Cre, Tfam LoxP/LoxP) and their littermate controls (DAT +/+, Tfam +/LoxP) were fed *ad libitum* and housed in standard conditions (constant temperature (22 ± 1°C), humidity (relative, 30%), and a 12-h light/dark cycle) approved and supervised by the Institutional Animal Care and Use Committee (IACUC) at ISU. Mice were weighed and subjected to behavioral tests every two weeks. Neurochemical, biochemical, and histological studies were performed following sacrifice at age 24 wks.

### Motor function test

For open-field test, a VersaMax system (VersaMax monitor, model RXYZCM-16, and analyzer, model VMAUSB, AccuScan, Columbus, OH) was used for monitoring locomotor activity. For horizontal and vertical activity and corresponding plots, mice were acclimated for 2 min prior to recording for 10 min using the VersaMax system. RotaRod equipment (AccuScan) was used to test coordination of movement as previously described ^77^. Briefly, time spent on rod rotating at 20 rpm was measured for a maximum of either 20 min or five trials, each of which ended with a mouse falling from the rod.

### Neuromuscular function and muscular strength

Each mouse was lifted over the grip strength meter (GSM)’s baseplate by the tail so that its forepaws could grasp onto the steel grip. Each mouse was then gently pulled backward by the tail until its grip released. The GSM measures the maximal force before the mouse releases the bar ^78^. Three trials were performed for each mouse with a 1-minute resting period between trials. Latency to release (sec) and gram-force (gF) were recorded.

### Social discrimination and novel scent tests

To determine the olfactory function of control and MitoPark mice, we used a social discrimination test as previously described ^79^. However, this procedure was adapted to use ANY-maze tracking software (AMS, Stoelting Co., Wood Dale, IL) to determine time spent sniffing based on the animal’s head being within a defined zone (1-cm perimeter around dish) surrounding the bedding. Total time spent sniffing the opposite sex’s bedding (from a group-housed cage) was recorded during a 3-min trial. Similarly, AMS was used to determine time sniffing a novel scent as described by Taylor *et al*. ^56^. During a 3-min trial, time spent sniffing scented and non-scented zones was recorded using AMS. Scents used were lemon, peppermint and vanilla, whereas water served as the non-scent.

### Cognitive testing

A six-day Morris water maze (MWM) protocol was used as described previously in an Alzheimer’s disease mouse model ^80^. Briefly, each mouse gets five 1-min trials per day. On the first day, the platform is visible and its position changes between trials to show that the ability to see and swim to the platform is not impaired by visual or motor deficits. On days 2-5, mice are placed into the MWM tank filled with white (Tempera paint), opaque water to learn to find a hidden platform whose position does not change between trials. The time required for a mouse to find and mount the platform is reported here as escape latency. Finally, the platform is removed on the sixth day when mice performed a single 1-min probe trial to show memory retention of the previously located platform, reported here as time spent searching in the quadrant that contained the platform during Days 2-5. Each trial was monitored using AMS. Each inter-trial interval was a minimum of 20 min, during which time mice were warmed and dried on heating pads placed under cages. Water temperature in the MWM tank was maintained at 23±1°C.

### Forced swim and tail suspension tests

For depressive phenotyping, tail suspension and forced swim tests were used measure behavioral despair or stress-coping response during inescapable tasks^81^. During tail suspension trials, mice were individually suspended at a height of 30 cm by attaching the tail to a horizontal ring stand bar using adhesive tape. Each 6-min test session was videotaped and scored using AMS for escape-oriented behavior/mobility and bouts of immobility. The time spent immobile was recorded for each mouse as a correlate of depression-like behavior ^8,82^.

For the Porsolt “forced swim” test, mice were placed individually in a glass cylinder (24 × 16 cm) with 15 cm of water maintained at 25°C as previously described ^83,84^. Mice were left in the cylinder and their behavior was videotaped from the side of the cylinder for 6 min. After the first 2 min, the total duration of time spent immobile was recorded during a 4-min test. A mouse was deemed immobile when it was floating 65% passively for at least 2.5 sec according to AMS.

### Elevated plus maze

Time spent in the open arms is inversely correlated with an anxious phenotype ^85^, which in rodents emerges as the trade-off between risk avoidance and spontaneous exploration of novel environments^86^. Mice were placed into the center of the elevated plus maze (Stoelting) and video-recorded for 10 min as previously described ^87^. Time spent in open arms was determined by AMS.

### Sleep latency test

Animals were allowed to acclimate 4 hours in the VersaMax monitor, and were then video-recorded from above after being awakened by gentle handling. Latency to sleep was determined by observer video-monitoring behavioral signs of sleep. Sleep was defined as 2 min of uninterrupted sleep behavior and 75% of the next ten minutes spent in sleep behavior as previously described ^56^.

### High performance liquid chromatography (HPLC)

Striatum, hippocampus, and olfactory bulb samples were prepared and processed for HPLC as described previously ^53^. Briefly, dissected brain regions were placed in a buffer comprising 0.2 M perchloric acid, 0.05% Na_2_EDTA, 0.1% Na_2_S_2_O_5_ and isoproterenol (internal standard) to extract monoamine neurotransmitters. Monoamine lysates were placed in a refrigerated automatic sampler (model WPS-3000TSL) until being separated isocratically by a reversed-phase C18 column with a flow rate of 0.6 mL/ min using a Dionex Ultimate 3000 HPLC system (pump ISO-3100SD, Thermo Scientific, Bannockburn, IL). Electrochemical detection was achieved using a CoulArray model 5600A coupled with a guard cell (model 5020) and an analytical cell (microdialysis cell 5014B) with cell potentials set at -350, 0, 150, and 220 mV. Data acquisition and analysis were performed using Chromeleon 7 and ESA CoulArray 3.10 HPLC Software and quantified data were normalized to wet tissue weight.

### Western blot

Protein lysates from the striatum and SN were prepared in RIPA buffer with protease and phosphatase inhibitors and ran on a 12-15% SDS-PAGE as previously described ^88^ before being transferred to a nitrocellulose membrane. After blocking for 1 h, membranes were incubated with primary antibodies at 4°C overnight. Next, membranes were incubated with secondary antibodies (Alexa Fluor 680 and Rockland IR800) at RT for one hour and images were captured via a LI-COR Odyssey imager. Densitometric analysis was done using ImageJ software.

### Statistical analysis

All behavioral tests were analyzed by two-way ANOVA with Bonferroni post-tests in GraphPad Prism software. Biochemical and neurochemical analyses were performed by two-tailed Student’s t-test. No data sets analyzed for this study violated the normality assumption, so nonparametric tests were not used. For ANOVA and t-tests, alpha level of p≤0.05 was used to determine significance. Plotted data represent mean +/- standard error of the mean.

## Data Availability

Raw data supporting the results reported in this article are in the figure source data files available upon request.

## Acknowledgements

This study was supported by NIH grants ES10586, NS074443 and NS039958. We would like to thank Gary Zenitsky for his help in preparing the manuscript.

## Contributions

M.R.L. conceived, designed and performed experiments, analyzed data and wrote the manuscript. S.G., M.A., and B.N.P. performed experiments. H.J. and V.A. provided intellectual input on experimental design, interpretation, and manuscript preparation. A.G.K. and A.K. led the investigation and helped conceive the project. All authors reviewed and edited the manuscript.

## Abbreviations

(αSyn): Alpha-synuclein
(AMS): ANY-maze software
(ANOVA): Analysis of variance
(CREB): cAMP response element-binding protein
(DG): dentate gyrus
(DAT): dopamine transporter
(DOPAC): 3-4-dihydroxyphenylacetic acid
(FST): forced swim test
(GI): gastrointestinal
(GSM): grip strength meter
(HPLC): high performance liquid chromatography
(HVA): homovanillic acid
(IHC): immunohistochemistry
(ISU): Iowa State University
(LRRK2): leucine-rich repeat kinase 2
(L-DOPA1): levodopa
(MPTP): 1-methyl-4-phenyl-1,2,3,6-tetrahydropyridine
(MWM): Morris water maze
(OB): olfactory bulb
(6-OHDA): 6-hydroxydopamine
(PD): Parkinson’s disease
(PINK1): PTEN-induced putative kinase 1
(qPCR): quantitative polymerase chain reaction
(RMS): rostral migratory stream
(SN): substantia nigra
(SVZ): subventricular zone
(SGZ): subgranular zone
(STR): striatum
(TST): tail suspension test
(TFAM): mitochondrial transcription factor A.

